# A Single Mutation in TRPC6 Protects Mice from Acute Lung Injury by Regenerating Endothelium

**DOI:** 10.1101/2025.02.19.639142

**Authors:** Mumtaz Anwar, Vijay Avin Balaji Raghunathrao, Ruhul Amin, Vigneshwaran Vellingiri, Jagdish Chandra Joshi, Md Zahid Akhter, Mohammad Tauseef, Ibrahim Hassan El-Erian, Nazanin Khaki, Ibra S. Fancher, Hazem Abdelkarim, Irena Levitan, Konstantinos Chronis, Vadim Gaponenko, Dolly Mehta

## Abstract

Regenerating vascular endothelium under sepsis, trauma, and viral infections is vital for promoting the resolution of inflammatory diseases such as acute lung injury (ALI). Transient receptor potential canonical (TRPC) channels mediated Ca^2+^ entry compromises organ functions and survival from lung injury. Through decoding the domain in TRPC6 responsible for vascular injury, we unveiled the intricate molecular mechanisms underlying vascular regeneration in injured tissue. We found that the substitution of isoleucine^111^ within the I^st^ ankyrin domain of TRPC6 for its isomer ^111^leucine (I^111^L-TRPC6) altered channel localization at the membrane, blocked TRPC6-mediated Ca^2+^ entry and cation currents without affecting TRPC6 protein expression. Next, we delivered WT-TRPC6 and I^111^L-TRPC6 to the endothelial cells (ECs) of TRPC6 knockout mice using liposomes and found that while WT-TRPC6 induced lung vascular inflammatory injury and EC death these responses were blocked in lungs expressing I^111^L-TRPC6 mutant. Instead, the I^111^L-TRPC6 mutant promoted lung EC proliferation and prevented vascular injury. These responses were recapitulated in a preclinical mouse model of ALI after injection of engineered TRPC6-blocking peptide, suggesting a novel strategy for regenerating anti-inflammatory vascular niche and preventing ALI therapeutically.

## Introduction

Acute lung injury and acute respiratory distress syndrome (ARDS) cause 40% mortality in affected patients and are characterized by protein-rich edema fluid and unchecked chronic inflammation (*1–3*). The lung endothelial cells (ECs) have long been the focus of ALI and ARDS research due to their ability to enhance vascular permeability and immune cell infiltration leading to inflammatory injury, and their propensity to resolve lung injury by launching a developmental program for restoring anti-inflammatory niches (*4–7*). Therapy that can enhance EC’s ability to restore the anti-inflammatory niche during injury is therefore pivotal in the treatment of ALI and ARDS, and in preventing death in hospitalized patients due to infections.

TRPC6 (transient receptor potential canonical) channel belonging to the TRPC superfamily is a transmembrane protein that mediates entry of divalent and nonselective cations (*8–11*). TRPC6, activated by DAG downstream of the ligand-receptor coupled pathway, is fundamental for providing cellular Ca^2+^ needed for signal transduction, gene transcription, and cellular structures and is ubiquitous in organisms across all phyla and kingdoms (*12–14*). In ECs, TRPC6 mediated Ca^2+^-entry in response to diverse agonists, including the endotoxin LPS (*15, 16*). Activated TRPC6 disrupted endothelial barrier function and enhanced inflammatory signaling by orchestrating interaction between MLCK and TLR4 pathway (*16–18*). Thus, mice lacking TRPC6 globally or in parenchyma were protected from ALI and polymicrobial- and LPS-induced sepsis (*16*). TRPC6-mediated Ca^2+^ influx is also crucial in regulating smooth muscle and glomerular podocyte function (*19–21*). Hyperactive TRPC6 has been linked to maladaptive tissue and organ remodeling, such as pulmonary hypertension (*13, 22, 23*). In addition, gain-of-function mutations in TRPC6 are associated with hereditary focal segmental glomerulosclerosis (FSGS), a renal disorder that can lead to end-stage renal disease (*24, 25*). Current evidence indicates sites for agonists and antagonists within the transmembrane domain of TRPC6 (*26*). However, a limited understanding of molecular mechanisms that modulate the activity of this channel hinders the development of potent and selective inhibitors of TRPC6.

TRPC6 contains 4-distinct ARD in the cytosolic portion of its amino terminus, which participate in protein-protein interactions and are essential for tetrameric channels and most mutations inducing TRPC6 activity in FSG were clustered within ankyrin domain 1 (G109S, P112Q, N143S). Here we show that substituting isoleucine^111^ within the I^st^ ARD for its isomer leucine^111^ blocked TRPC6-mediated increase in Ca^2+^ levels. Moreover, using NMR, we found that in contrast to WT, the I^111^L mutant induced allosteric transitions in TRPC6, impairing channel organization and cell-surface expression. LPS induced EC death in TRPC6 null mice expressing WT channel in EC but enhanced EC proliferation in mice expressing mutated TRPC6 channel (I^111^L-TRPC6) and these mice were protected from inflammatory vascular injury. Allosteric peptide developed against the TRPC6 I^111^L domain promoted EC proliferation dampening inflammation and vascular injury, defining the target for drug discovery against TRPC6 channelopathy.

## Results

### Characterization of the amino acid residues that can be targeted to block TRPC6 activity

TRPC6 contains 4 Ankyrin repeat domains (ARD), essential for channel formation (*27–30*). Point mutations in ankyrin domain 1 are linked with hyperactivation of channel and disease pathology(*25, 31–34*). To confirm that ankyrin 1 is required for channel expression, we transfected WT-TRPC6, TRPC6 lacking all ARD (ΔAnk-TRPC6), or TRPC6 lacking ankyrin 1 domain (ΔAnk1-TRPC6) in HEK cells, which minimally express TRPC6 (*35, 36*) (**Fig. 1Ai-ii**). As expected, we failed to find channel expression in cells expressing ΔAnk-TRPC6 (**Fig. S1A**).

**Figure 1:**
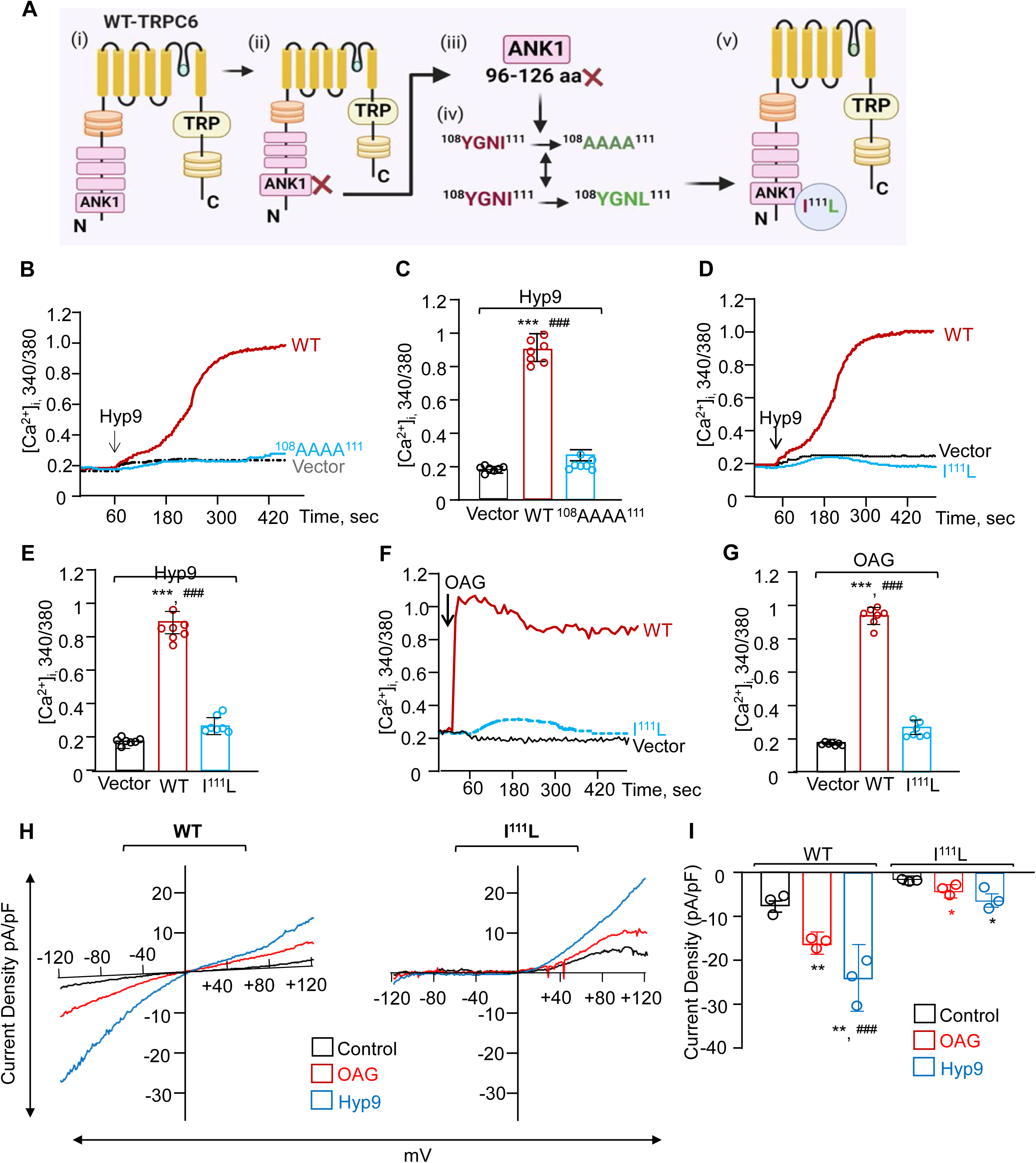
Functional analysis of I^111^ in regulating TRPC6 activity. **(A)** Schematics of WT-TRPC6 channel and identification of residues required for channel activity. (**B-G**) Ratiometric analysis of intracellular Ca^2+^ in HEK cells expressing WT-TRPC6 or TRPC6 mutant after addition of 1 µM Hyperforin 9 (Hyp9; **B-E**) or 100 µM OAG (**F-G**). **B**, **D**, and **F** shows a representative tracing of the average response of 10–20 cells while **C**, **E**, and **G** shows mean ± S.D. of increased intracellular Ca^2+^ calculated as fold. Experiments were repeated three times. (**H-I**) Current tracings in HEK cells expressing WT-TRPC6 or TRPC6 mutant obtained during voltage ramps from −120 mV to + 120 mV in response to Hyp9 and OAG. Currents are leak-subtracted. **I**, shows mean peak current densities of Hyp9 and OAG-elicited currents obtained at a holding potential of −120 mV from 3 dishes expressing WT or I^111^L mutant. All data in **C, E, G and I** are mean ± SD. One-way ANOVA with Tukey’s post hoc. **p < 0.001, ***p < 0.0001 compared to vector. ##p < 0.001, ###p < 0.0001 compared to mutants. See also Supplement Figure 1.

Compared to WT-expressing cells, expressing ΔAnk1-TRPC6 showed 35% expression of TRPC6 **(Fig. S1A).** Next, we deleted residues 96-111 (ΔAnk1A-TRPC6) in the TRPC6 ankyrin-1 domain (**Fig. 1Aiii**). We stimulated these cells with Hyp9, an activator of TRPC6 (*37*), to assess the impact of deletion of these residues on channel assembly and Ca^2+^ entry. Our rationale was that residue P to Q mutation at residue 112 hyperactivated the channel (*25, 31, 38*). Interestingly, compared to WT-TRPC6 expressing cells, Hyp9-induced Ca^2+^ entry was reduced by 80% in cells expressing ΔAnk1-TRPC6 or ΔAnk1A-TRPC6 mutant (**Fig. S1B-C**). Basal cytosolic Ca^2+^ concentration did not differ between cells expressing the vector, versus WT-TRPC6, ΔAnk1-TRPC6, or ΔAnk1A-TRPC6 mutants (data not shown). ΔAnk1A-TRPC6 mutant expression was also like ΔAnk1-TRPC6 at the protein level, indicating that amino acids 96-111 are required for channel formation and function (**Fig. S1D**).

To find the residues with 96-111 that may block channel activity, we performed structural analysis using the ELM database, which predicted YGNI [aa108-aa111] as the motif likely regulating channel organization and binding with Ca^2+^. Therefore, we mutated these ^108^YGNI^111^ to ^108^AAAA^111^ in TRPC6 and determined cytosolic Ca^2+^ concentration and channel expression. Hyp9 failed to induce Ca^2+^ entry in HEK cells, traducing ^108^AAAA^111^ TRPC6 mutant (**Fig.1 B-C**). The mutant was still expressed around 50% of the WT channel (**Fig. S1E**), indicating that targeting the YGNI motif may identify the residue critical for channel formation and activity.

Further structural analysis of TRPC6 suggested that slight perturbation at I^111^position can significantly affect TRPC6 organization. We tested this hypothesis by substituting I^111^ with A or its isomer leucine. Both mutants were similarly expressed as WT-TRPC6 (**Fig. S1F-G**). Importantly, substituting I^111^ to L^111^ residue in TRPC6 completely blocked Ca^2+^ entry (**Fig. 1D-E**). Next, we stimulated HEK cells by transducing WT or I^111^L-TRPC6 mutants with OAG, a cell-permeable analog of DAG produced during receptor stimulation (*18, 39*). I^111^L-TRPC6 mutant also compromised OAG-induced Ca^2+^ entry (**Fig. 1F-G**).

To validate the role of this residue in regulating cationic current, we performed whole-cell patch-clamp recordings in HEK cells transducing the WT-TRPC6 or the I^111^L-TRPC6 mutant. After adding OAG and Hyp 9, we observed a cationic current in cells transduced with WT-TRPC6, with the peak response occurring within 2-5 min compared to cells transduced with I^111^L-TRPC6 (**Fig. 1H**). Hyp9 and OAG produced similar outward currents at +120 mV in WT-TRPC6 expressing cells and cells transducing I^111^L-TRPC6, indicating that nonselective cation channels mediated these currents. (**Fig. 1H**). However, inward currents at −120 mV amplitude were inhibited by more than 90% in cells transducing I^111^L-TRPC6 mutant than in WT-TRPC6 transducing cells (**Fig. 1H-I)**. These results identified isoleucine I^111^ as the crucial residue indispensable for TRPC6-mediated Ca^2+^ entry.

### I^111^L-TRPC6 impairs TRPC6 organization and surface localization

We next assessed receptor localization by depleting TRPC6 in ECs using shRNA and expressing WT-TRPC6 or I^111^L-TRPC6 mutant in TRPC6-depleted ECs **(Fig. 2A)**. Using total internal reflection fluorescence microscopy (TIRFM), which limits fluorescence detection up to 100 nm and thus detects plasma membrane localization (*40, 41*), we observed that WT-TRPC6 was expressed at the cell surface. However, 80% of I^111^l-TRPC6 was internalized (**Fig. 2B-C**). We further validated these findings biochemically using surface biotinylation assay, which also showed that WT-TRPC6 was expressed on the cell surface as compared to the mutant (**Fig. 2D-E**).

**Figure 2.**
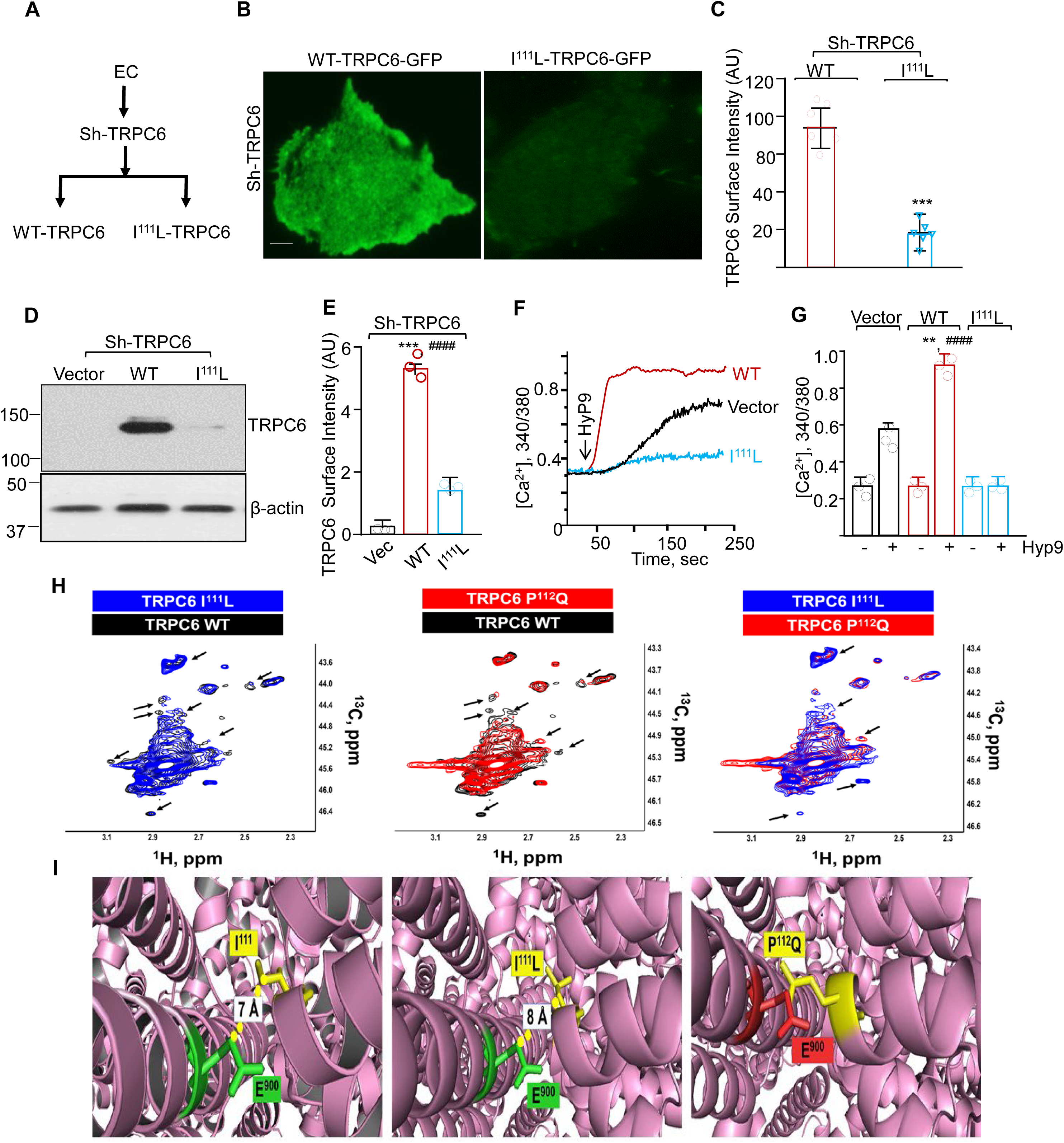
Cell surface localization and organization of WT and I^111^L-TRPC6 mutant. **(A)** Schematics. (**B-C**) WT or I^111^L TRPC6 expressing ECs visualized using TIRF microscopy. **B** shows a representative image while **C** shows mean pixel intensity (line) at the cell surface. N=3. Scale bars: 20 µm. (**D-E**) Cell surface TRPC6 assessed after biotinylation followed by immunoprecipitation (IP) with anti-TRPC6 antibody. Total cell lysates were assessed for actin to confirm equal protein loading. **D** shows a representative blot while **E** shows the mean ± SD fold change in biotinylated TRPC6 versus total actin expression. Experiments were repeated multiple times. (**F**-**G**), Human lung EC transducing indicated cDNAs were stimulated with HyP9 and [Ca^2+^]_I_ was determined as in Fig. 1. **F** shows a representative mean calcium response (8-10 cells/group) from two independent experiments while **G** shows mean ± S.D. of increase in intracellular Ca^2+^ calculated as fold increase compared to vector with or without Hyp9 stimulation. (**H-I**), NMR showing conformational differences in cells transducing WT, I^111^L and P^112^Q TRPC6 channel. The site of the I^111^L mutation in the ARD is shown as a red circle in the proximity of the pole helix. The more compact size of leucine allows the ankyrin repeat domain to closely approach the pole helix, as indicated by dark arrows. This movement can trigger the repositioning of other domains (shown with light grey arrows) and eventually alter the relative positions of pore-forming helices 5 and 6 and change ligand binding (**I**). All data in **C, E,** and **G** are shown as mean ± SD). **C**, Paired two tailed t-test. ***p < 0.0001 compared WT. **E and G,** One-way ANOVA with Tukey’s post hoc. *p < 0.05, ***p < 0.0001 compared to vector. ###p < 0.0001 compared to I^111^L. See also Supplement Figure 2.

Based on these above studies, we addressed the possibility that I^111^L-TRPC6 mutant will block intracellular Ca^2+^ ([Ca^2+^]_i)_ increase in response to Hyp9 and thrombin. Thrombin generates DAG (Nakashima S, 1991). TRPC6-depleted ECs transducing WT or I^111^L-TRPC6 were loaded with the Ca^2+^-sensitive dye, Fura2, and stimulated with Hyp9. Compared to the vector expressing EC, Hyp9 increased [Ca^2+^]_i_ in ECs transducing WT-TRPC6 but modestly in cells transducing the I^111^L-TRPC6 (**Fig. 2F-G**). In other studies, these ECs were co-transfected with a Ca^2+^ sensor, pcDNA-RCaMP (*42*) after which we monitored Ca^2+^ transients using thrombin. Thrombin rapidly increased pcDNA-RCaMP expression in ECs expressing WT channel but not in ECs expressing I^111^L-TRPC6 mutant **(Fig. S2, A, and B)**. These findings demonstrate that internalized I^111^L-TRPC6 fails to induce intracellular Ca^2+^ transients.

A rise in intracellular calcium is crucial in disrupting endothelial barrier function (*43, 44*). Thus, we determined endothelial permeability by measuring trans-endothelial electrical resistance (TEER) in ECs expressing WT-TRPC6 and I^111^L-TRPC6 in response to thrombin. We found that expression of WT-TRPC6 decreased TEER, which was significantly reduced in ECs expressing I^111^L-TRPC6 mutant (**Fig. S2, C-D**).

We then assessed the structural effects of the I^111^L mutation by NMR on reductively methylated TRPC6 and compared them with reductive methylated WT and P^112^Q mutant (**Fig. 2H**). After confirming that the signals from TRPC6 were dominant in the recorded spectra (data not shown), we assessed spectral comparison, which revealed multiple changes in chemical shifts and linewidths in WT, I^111^L, and P^112^Q TRPC6 (**Fig. 2I**), indicating that these point mutations cause TRPC6 to assume a conformation that differs from that of the WT channel. Intriguingly, I^111^L- and P^112^Q-TRPC6 also had distinct conformations, apparently correlating with the observed differences in the effects of these mutations on the function of TRPC6. This observation agrees with the agonist and antagonist-induced long-range allosteric transitions that converge at ARD1 of TRPC6 (*26*).

### EC-specific expression of I^111^L-TRPC6 mitigates lung vascular hyper-permeability

We have shown that LPS generates DAG, which requires TRPC6 and TLR4 expression on EC to induce Ca^2+^ entry and lung injury (*16*). We, therefore, assessed if EC-specific expression of I^111^L-TRPC6 is sufficient to impair lung hyperpermeability *in vivo*. We complexed WT- or I^111^L-TRPC6 cDNA driven by the Cdh5 promoter (to restrict gene expression only in EC) into liposomes using established techniques and delivered them *i.v.* into *TRPC6^-/-^* mice (**Fig. 3A**). At 48h post-transfection, we confirmed using imaging and western blotting that WT and mutated channels were expressed in ECs (**Fig. S3 A-C**). We next challenged these mice with *i.t.* LPS (2.5 mg/kg) and found that LPS increased inflammation to a similar level at 4 and 16 h (**Fig. S3D**). We, therefore, focused on assessing the role of WT versus mutated TRPC6 channel on edema formation at 4 and 16h. Restoring WT-TRPC6 in EC rescued edema formation at 4 and 16h after the LPS challenge in TRPC6^-/-^ mice (**Fig. S3E and Fig. 3B**). However, LPS failed to increase edema formation in TRPC6^-/-^ mice transducing I^111^L-TRPC6 in EC (**Fig. 3B and S3E**). Intriguingly, EC-restricted expression of I^111^L-TRPC6 suppressed lung injury as revealed by decreased septal thickening, leukocyte infiltration, and inflammatory score (**Fig. 3C-D**). MPO staining of these lung sections validated the results of inflammation and injury (**Fig. 3E-F**)

**Figure 3.**
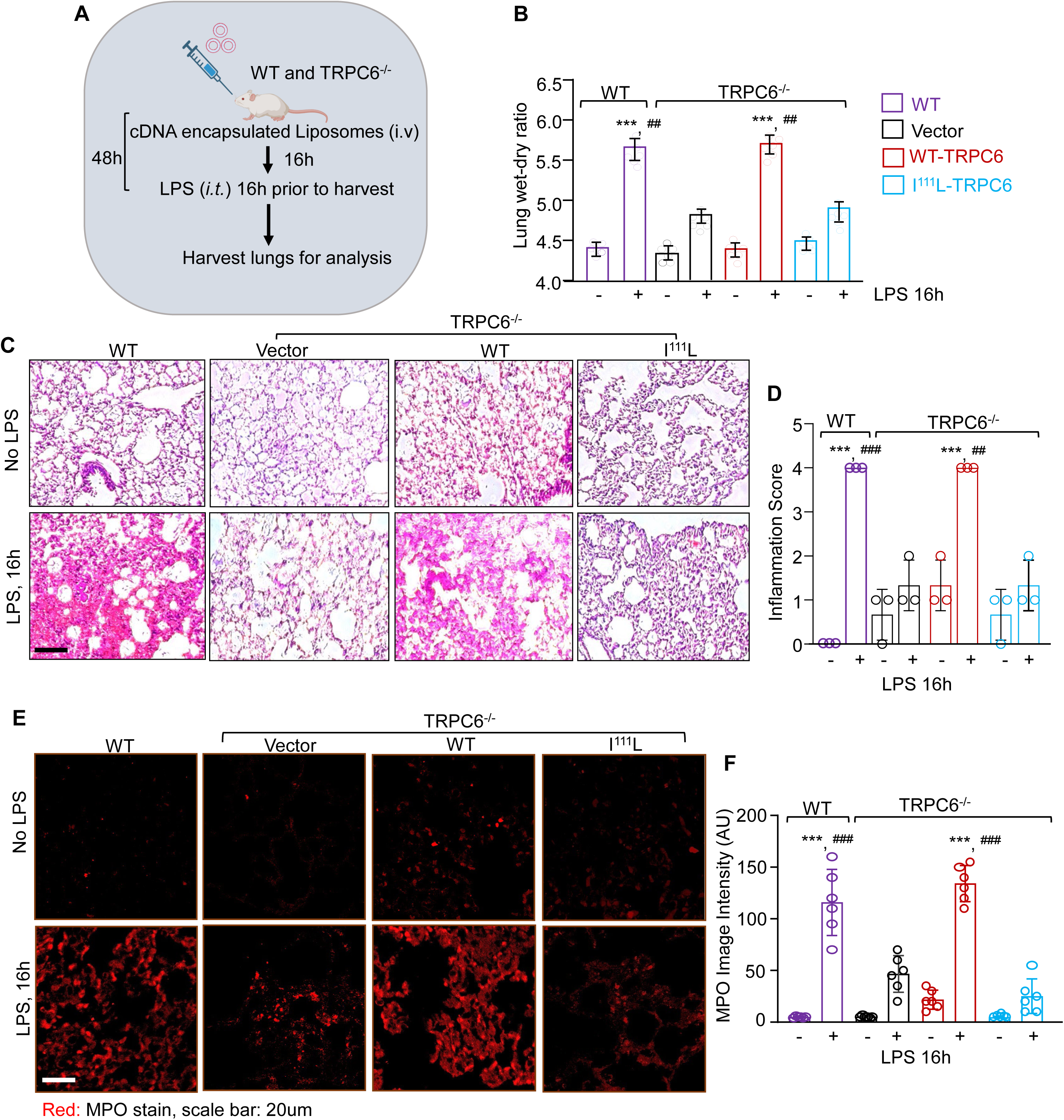
I^111^L-TRPC6 mutant expression exclusively in endothelium suppress lung inflammatory injury. **(A)** Schematics. **(B-D)** edema, alveolar septal thickening, leukocyte infiltration, and inflammation in WT or *Trpc6^−/−^* mice expressing vector, WT- or I^111^L TRPC6. (**E-F)** MPO intensity expressed as arbitrary unit (A.U) in **F**. **B, D,** and **F,** One-way ANOVA with Tukey’s post hoc. ***p < 0.0001 compared to no LPS. ##p < 0.001, ###p < 0.0001 compared to LPS receiving mice. All data in **B, D,** and **F** are shown as mean ± SD). See also Supplement Figure 3.

### Mutated-TRPC6 promotes EC regeneration during lung injury

EC death is crucial in augmenting lung vascular inflammatory injury (*7, 45–47*). We, therefore, tested the hypothesis if TRPC6 activity favors endothelial cell regeneration over cell death during injury resolving lung inflammatory injury. Using Ki67 and annexin, we first assessed endothelial cells proliferation and death in ECs expressing WT or I^111^L-TRPC6 mutant following LPS exposure. We found that ECs expressing I^111^L-TRPC6 were proliferative basally and resisted death from LPS. In contrast, ECs expressing WT did not proliferate and died after LPS challenge **(Fig. S4A-D)**. Next, we assessed ECs regeneration *in vivo* in WT and *Trpc6^-/-^* mouse lungs traducing vector, WT or mutated channel (**Fig. 4A**). We found no statistical difference in EC proliferation between WT and *Trpc6^-/-^* mouse lungs traducing vector, WT or mutated channel at baseline (**Fig. 4B-C**). However, after injury, *Trpc6^-/-^* lungs expressing mutated channel (I^111^L-TRPC6 mutant) showed markedly increased EC proliferation than TRPC6 null mice expressing vector alone or WT channel (**Fig. 4B-C)**. In other studies, we administered BrdU in WT and *Trpc6^-/-^* mouse lungs, transducing the above-mentioned constructs after challenging the mouse with LPS. Likewise, we found increased BrdU+ cells in *Trpc6* null lungs transducing I^111^L-TRPC6 vector or I-L mutant (**Fig. 4D).** These mice also showed increased EC death in *Trpc6* null mice receiving WT channel than vector or mutated channel (**Fig. 4E-F**). These findings demonstrate that loss of channel activity promotes EC proliferation during injury.

**Figure 4.**
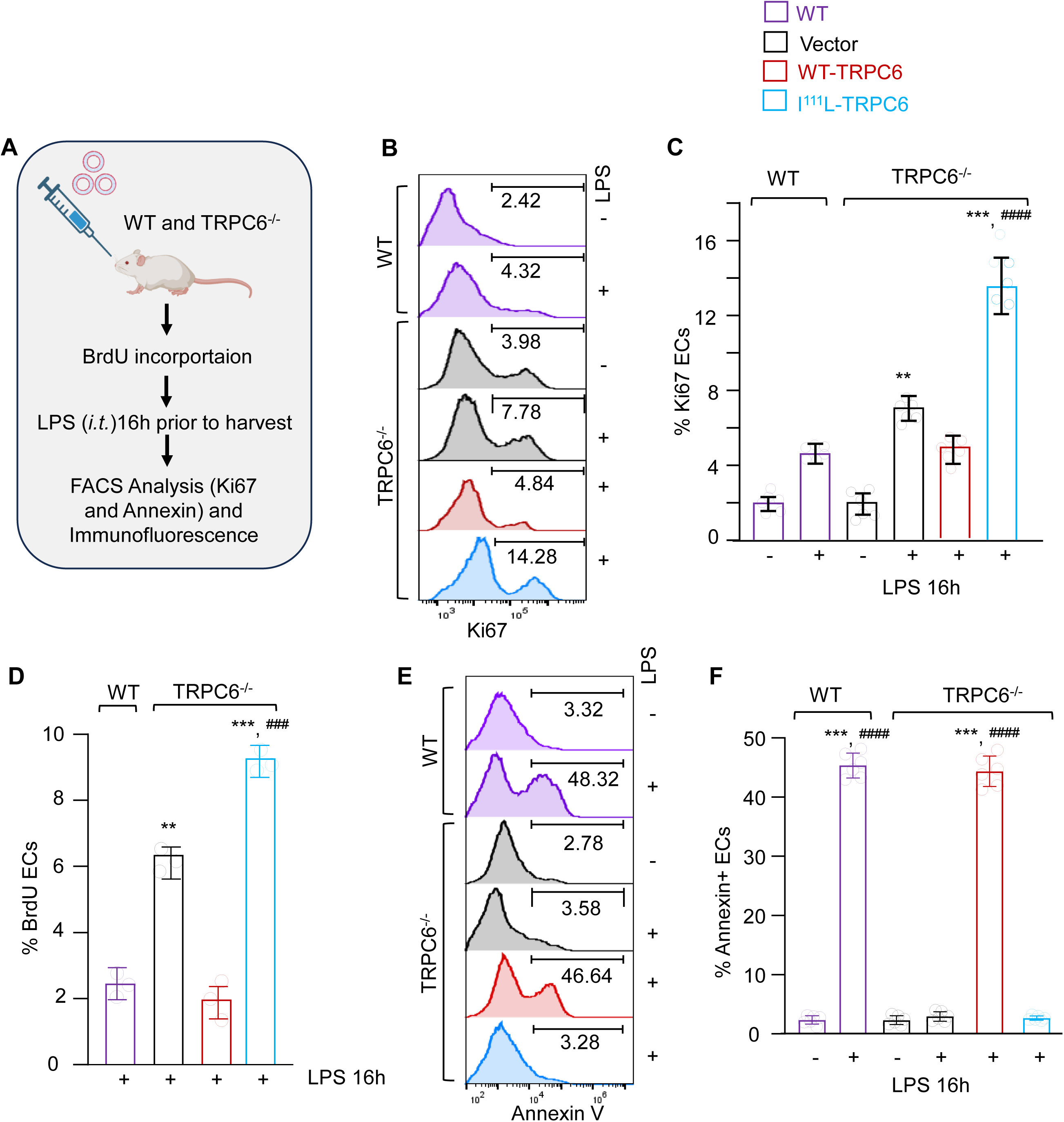
I^111^L-TRPC6 mutant promotes endothelial proliferation. **(A)** Schematics. (**B-C**) Histogram (**B**) and quantitation (**C**) of EC proliferation in single lung suspension from WT or *Trpc6^−/−^* lungs expressing vector, WT- or I^111^L using anti-Ki67 antibody. (**D**) Quantitation of BrdU expression in EC as in B-C. (**E-F**) Histogram (**E**) and quantitation (**F**) of EC death using anti-annexin V antibody. **C, D, F** One-way ANOVA with Tukey’s post hoc. **p < 0.001, ***p < 0.0001 compared to unstimulated. ###p < 0.0001 compared to LPS receiving lungs All data in **C, D,** and **F** are shown as mean ± SD. See also Supplement Figure S4

### TRPC6 blocking peptide promotes EC proliferation and prevents lung inflammatory injury

Based on the above findings, we explore the effect of inhibiting TRPC6 channel activity by designing a peptide based on the TRPC6 ankyrin 1 motif (Myr-QDISSLRYE), which we named TRPC6-BP. We pretreated ECs with control peptide (Seq) and TRPC6-BP for an h and then loaded them with Flou4 to determine changes in intracellular Ca^2+^calcium levels. TRPC6-BP inhibited thrombin-induced increase in Ca^2+^ levels (**Fig. S5 A-B**). Because loss of TRPC6 inhibited LPS-induced Ca^2+^ entry (*16*), we also assess the effect of TRPC6-BP on LPS-induced increase in Ca^2+^ levels. Strikingly, TRPC6-BP also inhibited LPS-induced increase in Ca^2+^ levels (**Fig. S5 C-D**). We next used a mouse model of ALI to determine the *in vivo* effects of TP6 in inducing EC proliferation, inhibiting vascular injury and inflammation. We administered TP6 in WT mice before (Preventive study, **Fig S6A**) or after *i.t* LPS injection (Therapeutic study, **Fig. 5A**). In both models, TP6 promoted EC proliferation despite injury and prevented EC apoptosis (**Fig. 5B-E and Fig. S6B-E**). Interestingly, TP6 effectively inhibited ALI as reflected by markedly decreased lung edema and cytokine expression (**Fig 5F-G and Fig. S5F-G**). Taken together, these data suggest that TP6 attenuates vascular hyperpermeability and inflammatory injury in mice model of ALI through rejuvenating endothelium, indicating its potential effectiveness in preventing ALI.

**Figure 5.**
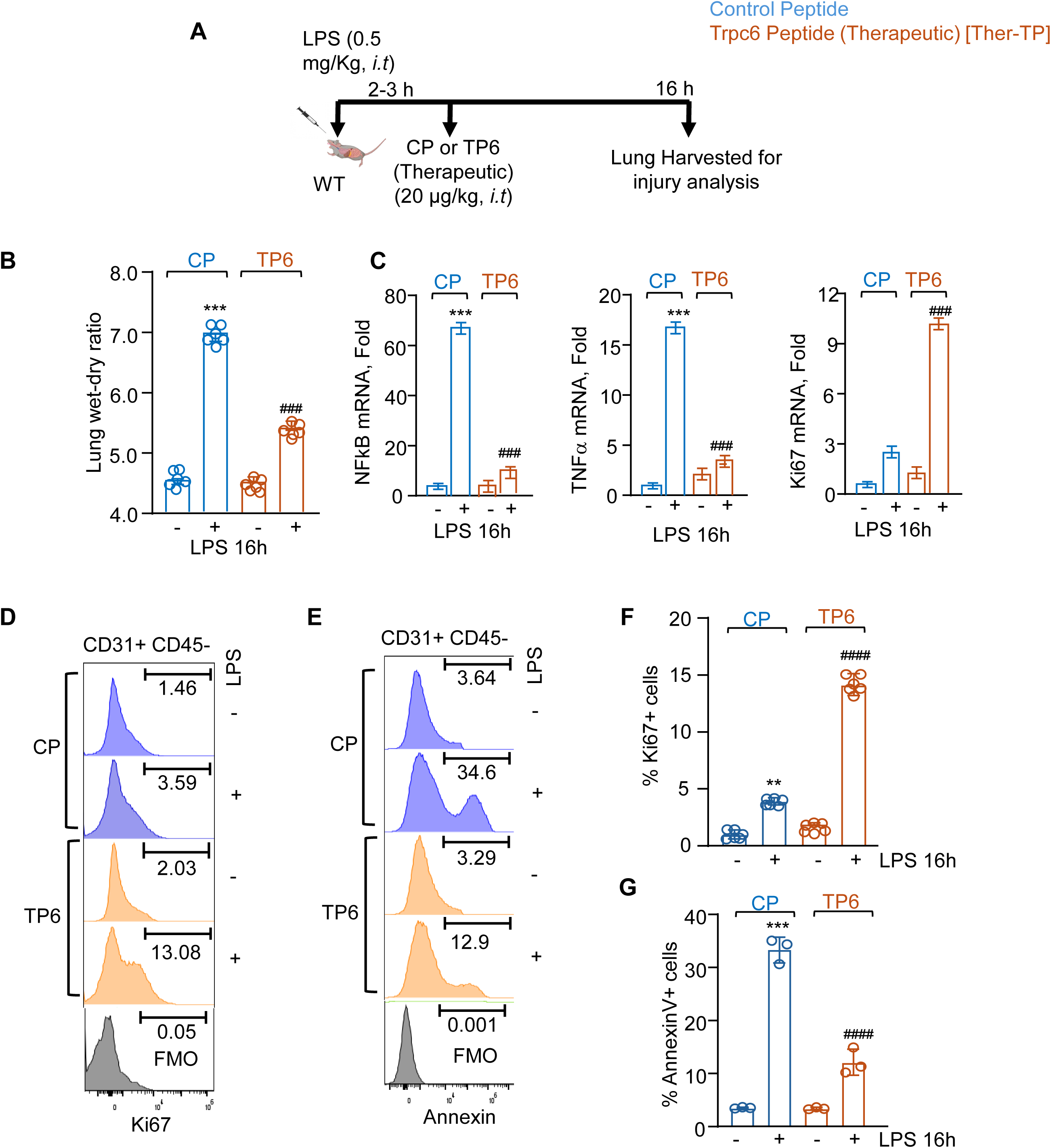
TRPC6 blocking peptide promotes endothelial cell proliferation and suppresses LPS-induced lung injury and inflammation. (**A**) Schematics of peptide delivery. (**B-E**) proliferation and death using Ki67 and annexin V antibodies in EC (CD31+ CD45-). **B-C** show histogram while **D-E** show quantitation. (**F**) Lung wet/dry ratio. N =6 mice per group. (**G**) Gene expression normalized against GAPDH. N=3. One-way ANOVA with Tukey post hoc test was used in **D** and **E** while one-way ANOVA with Fisher least significant difference test was used in **F** and **G**. See also Supplement Figure S5-6. **p, < 0.001***p < 0.0001 compared to vehicle. ###p < 0.0001 compared to LPS.

## Discussion

In this study, we demonstrate the critical role of I^111^ residue in mediating Ca^2+^ entry and cationic currents, the mutation of which effectively rescues the anti-inflammatory EC niche, resolving vascular inflammatory injury during ALI. We further show that the TRPC6-blocking peptide, TRPC6-BP, raised against this domain inhibits the TRPC6 activity in ECs, promotes their proliferation and effectively inhibits inflammation resolving ALI. The protective effect of TRPC6-BP was shown even when the treatment was started after instilling LPS. Thus, our study provides strong support for the concept of simultaneously promoting EC proliferation while reducing vascular leakage and maintaining lung-fluid homeostasis as an effective strategy for treating ALI. The TRPC6 was initially discovered as a DAG-sensitive calcium channel (*10, 48*). TRPC6 contains 6 transmembrane domains with 4 ankyrin-like repeat domains (ARD) at the N-terminus, a pore region located between domains 5 and 6, 2 coiled-coil domains (CC), and a conserved TRP box at the C-terminus (*27, 29, 49*). ARDs are required for channel formation and thereby channel activity (*50–53*). We showed that TRPC6 expression was reduced by 70% at the protein level in cells expressing the TRPC6-ΔAnk1 or TRPC6-ΔAnk1A mutants and importantly deletion of amino acids 96 to 111 from the I^st^ ARD (TRPC6-ΔAnk1A) demonstrated that these amino acids were required for channel formation and TRPC6 activity. High-resolution cryo-electron microscopy has identified novel modulation sites for agonists and antagonists within the transmembrane domain of TRPC6 (*26*). Moreover, gain of function mutations in the TRPC6 channel such as P^112^Q have been identified in patients suffering from focal and segmental glomerulosclerosis(*25*). We showed that Hyp9 and OAG failed to induce Ca^2+^ entry in cells transducing the I^111^A or I^111^L-TRPC6 mutant without altering TRPC6 expression. Moreover, the OAG and Hyp9-induced inward currents were inhibited by greater than 90% in HEK cells transducing the I^111^L-TRPC6 mutant.

Studies show that TRPC6 is translocated from intracellular compartments to the plasma membrane upon ligand stimulation(*38*). We showed that WT-TRPC6 was expressed at a higher level at the plasma membrane than the mutant. P^112^Q mutation induces TRPC6 activity by altering its conformation (*24, 54*). Spectral comparison revealed multiple changes in chemical shifts and linewidths in WT, I^111^L, and P^112^Q TRPC6 indicating that these point mutations cause TRPC6 to assume a confirmation that differs from that of the WT channel. Intriguingly, I^111^L- and P^112^Q-TRPC6 also had distinct conformations, apparently correlating with the observed differences in the effects of these mutations on the function of TRPC6.

ECs massively die during inflammatory injury (*55–57*). We showed that while WT-TRPC6 augmented EC death the mutant promoted EC proliferation Because activated TRPC6 induces RhoA and MLCK pathway leading to EC contraction, vascular hyperpermeability, and TLR4 signaling we infer that WT-TRPC6 induces hyperpermeability and inflammatory injury. Thus, TRPC6 deficiency prevented TLR4-mediated signaling in EC rendering TRPC6^-/-^ mice resistant to endotoxin- and sepsis-induced lung injury (*16, 21*). Here, we showed that a single point mutation within TRPC6 (I^111^L-TRPC6) was sufficient to reduce its ability to disrupt junctions. We also showed that mutant blocked vascular injury and instead induced proliferation of ECs.

There are no specific pharmaceutical tools at present to downregulate TRPC6 functions in the setting of injury because of non-specific effects of small molecule inhibitors of TRPC6. Moreover, the mechanisms by which these molecules inhibit TRPC6 activity remain unclear.

## Supplementary Figure Legends

**Figure S1: Identification of amino acids regulating TRPC6 activity. (A and B)** protein expression in HEK cells expressing WT-TRPC6 (WT) or TRPC6 mutants (ΔAnk, ΔAnk1) using anti-TRPC6 (top) or anti-actin (bottom). Numbers in parenthesis show densitometric analysis from experiments that were performed three times independently. **(C and D)**, Intracellular Ca^2+^ in response to 1 µM Hyperforin 9 (Hyp9) in HEK cells expressing indicated cDNA after loading with Fura 2-AM for 15 min. A representative tracing shown in **C** is the average response of 10–20 cells while the **D** shows mean ± S.D. of intracellular Ca^2+^ increase calculated as fold over empty vector n=5 coverslip/group that were assessed independently two times. (**E-G**), TRPC6 expression in HEK cells expressing indicated cDNAs taking actin (bottom) as the loading control. Data are expressed as mean ± SD. Numbers in parenthesis show densitometric analysis from experiments that were performed three times independently. One-way ANOVA with Tukey’s post hoc. ***p < 0.0001 compared to Hyp9. ###p < 0.0001 compared to mutant expressing and Hyp9 stimulated cells. Data in **D** shows as mean ± SD.

**Figure S2: I^111^L TRPC6 mutant inhibits cytosolic Ca^2+^ transients and barrier dysfunction.** (**A-B**) Time-lapse images (**A**) and quantification (**B**) of WT or mutant after photoconversion at time 0 (488 nm, green) and 50 nM thrombin addition (561 nm, red). 488 nm (green). Scale bar, 10 μm. N= 5–6 cells from experiments repeated two times. (**C-D**) TEER in real time was recorded in response to 50 nM thrombin in ECs expressing indicated cDNAs. **C** shows a representative TEER trace while **D** shows the quantification of the TEER from experiments that were repeated at least two times. One-way ANOVA with Tukey’s post hoc. **P < 0.001,***P < 0.0001 compared to thrombin. ###, P < 0.0001 compared with I^111^L TRPC6 without and with thrombin stimulation in **B**. All data in **B and D** are shown as mean ± SD.

**Figure S3: Restoration of TRPC6 expression in *Trpc6^−/−^* mice restore LPS-induced lung vascular inflammatory injury.** (**A**), Expression of WT and I^111^L TRPC6 in TRPC6 null lungs detected using anti-GFP antibody. (**B-C**) Lung cytokine expression (**B**) and edema (**C**) in WT and TRPC6 null mice expressing indicated cDNAs after receiving LPS. One-way ANOVA with Tukey’s post hoc. ***, P < 0.0001 compared to LPS. ### P < 0.0001 compared to I^111^L TRPC6 after LPS. All data in **B** and **C** are shown as mean ± SD.

**Figure S4. I^111^L-TRPC6 promotes endothelial proliferation.** (**A**) Schematics. (**B-D**) Histogram (B) and quantitation (**C-D**) of EC proliferation and death TRPC6 null ECs expressing indicated cDNAs using anti-Ki67 and anti-annexin V antibodies. **C, D** One-way ANOVA with Tukey’s post hoc. **p < 0.001, ***p < 0.0001 compared to unstimulated. ###p < 0.0001 compared to LPS. All data in **C** and **D** are shown as mean ± SD.

**Figure S5**: **Effect of TRPC6-blocking peptide in regulating Ca^2+^ transients**. **(A-D)** Intracellular Ca^2+^ in ECs in response to thrombin **(A-B)** or LPS **(C-D)** after 1.5 h treatment with 1 µM Myr-CP or the Myr-TRPC6-blocking peptides (TRPC6-BP). **A** and **C** show the representative images while **B** and **D** shows the quantification. One-way ANOVA with Tukey’s post hoc. ***, P < 0.0001 compared to 0 min. ###, P < 0.0001 compared to TP6. Experiments were repeated at least two times. All data in **B** and **D** are shown as mean ± SD. (n=3).

**Figure S6: Effect of TRPC6-blocking peptide in regulating EC proliferation and vascular injury in a preventive mice model of ALI.** (**A**) Schematic. (**B-C**) ECs proliferation (**B and D**) and EC death (**C and E**). Representative FACS plot (**B-C**) and quantitation (**D-E**). (**F-G**) Lung wet/dry ratio, n=6 (**F**) and cytokine expression (**G**) (n=3) in the mice pretreated with 20 μg/kg BW Myr-control or Myr-TRPC6 2 hours before LPS administration (1mg/kg i.t. BW). (C) mRNA profile of inflammatory and proliferative genes. Statistics: One-way ANOVA with Fishers least significant difference test in B and C. ***p < 0.0001 compared to LPS Unstimulated WT lungs treated with CP and TP6; ###p < 0.0001 compared to LPS stimulated WT lungs receiving CP in **B** and **C**. **p < 0.001 and ***p < 0.0001 compared to LPS Un-stimulated WT lungs treated with CP and TP6 in **F** and **G**. ###p < 0.0001 compared to LPS stimulated WT lungs treated with CP in **F** and **G**.

## Materials and Methods

### Cell culture

Human embryonic kidney cells (HEKs), human umbilical cord vein endothelial cells (HUVECs), and human pulmonary arterial endothelial cells (HPAEC) obtained from Lonza Allendale, NJ, USA, were used in all cell studies. Cells were cultured in a humidified atmosphere with 5% CO_2_ at 37°C in an EGM-2 medium supplemented with growth factors obtained from Lonza, 10% FBS, and penicillin-streptomycin antibiotics. All studies were conducted on HUVEC and HPAEC between passages 5-7 (*40, 58*).

### Animals

Mice used in this study were approved by the Institutional Animal Care and Use Committee of the University of Illinois at Chicago. All experiments were performed on 6–8-week-old mice weighing 20-25 grams. Sex-matched groups of male and female mice were used for these studies. No animals were excluded from the analysis. We calculated the sample size using software G Power (or G Power software) based on the pre-designed effect size between the groups based on Cohen’s principles at a power =0.80 and significance level = 0.05 (*59*).

### Cell culture and transfection

Cells were transfected at passages 5-7 using Santa Cruz transfection reagent or Amaxa electroporation. TRPC6 was depleted using shRNA 5’-AAACCAUCUUCAUCUUCCCUU -3’ (Genscript). Control shRNA (shCtr) ON-TARGETplus non-targeting pool, Genscript, was used as a control in all experiments. For transfection experiments, 1.5µg of shRNA (double knockdown) was used for 48 h. For transfecting cDNA, ECs were first transfected with shRNA for 48 h and then transduced with 2µg of cDNA for another 24 h. cDNA was transduced in depleted cells using FuGENE® HD transfection reagent (Promega #E2311) (*60*).

### Cell and tissue imaging

ECs were rinsed three times with Ca^2+^ and Mg^2+^ containing PBS and fixed with 2% paraformaldehyde. Cells were then incubated with 5% normal goat serum supplemented with 1% Bovine serum albumin in TBS (0.2M Tris base, 1.5M NaCl) for 2 h at room temperature. After rinsing three times with PBS, cells were incubated with 1:50 anti-CD31, anti-vWF (Santa Cruz) or anti-TRPC6 (Santa Cruz and Alomone) antibody for 1 h at room temperature, following which cells were rinsed again and incubated with 1:250 dilution of Alexa-Fluor 594 secondary Donkey anti-rabbit antibody (Thermo Fisher Scientific) at room temperature for 1h. Isotype control primary antibodies (Thermo Fisher Scientific) were used as a negative control to validate the antibodies’ specificity and eliminate the background signal. Formalin-fixed, 4-µm-thick lung sections were cut followed by subsequent histological analysis and MPO staining. Lung sections were visualized with light microscope.

### TIRF microscopy

TIRF microscopy was performed using a motorized laser TIRF imaging system (Carl Zeiss) equipped with an ORCA-Flash4.0 V3 Digital CMOS camera (Hamamatsu), and an α Plan-Apochromat 100×/1.46 NA objective (Carl Zeiss). For detection of cell surface expression of the receptor, GFP-tagged WT or TRPC6 mutant were imaged along with mCherry-tagged stargazin (cell surface marker) at λ = 488 nm and λ = 561 nm excitation for GFP and mCherry, respectively. Images from green and red channels (excitation 488 nm and 561 nm) were obtained by fast switching the excitation lasers using AxioVision software (Carl Zeiss).

### Image analysis

ImageJ-NIH was used to design the protocols for quantifying all confocal images using standard methods. An area of uniform size was used in different conditions to quantify fluorescence intensity and gap area between two endothelial cells. Moreover, mean immunostaining intensities (gray values of stained cells in 12-bit Zeiss CZI images) were measured.

### Fluorescence-activated cell sorting

Lungs were minced and digested with 1mg/ml collagenase A (Roche) for 30 min at 37°C, after which digested tissue was passed through a 75μm cell strainer to obtain single-cell suspensions. Cells were stained with anti-CD31 and anti-CD45 antibodies, and endothelial cells (CD45-CD31+) were sorted using the Beckman Coulter cell sorter as described (*45*).

### Quantitative real-time PCR

Total RNA was isolated using Trizol reagent (ThermoFisher Scientific) and quantified using BioDrop DUO+ (Biochrom, UK). RNA (1µg) was reverse transcribed using High-Capacity RNA to cDNA Kit (Applied Biosystems) according to the manufacturer’s protocol. The cDNA products were assessed using quantitative Real-Time PCR analysis and Fast SYBR™ Green Master Mix (Applied Biosystems). Each measurement was duplicated using a CFX384 real-time on Applied Biosystems QuantStudio Real-Time PCR System. The PCR conditions were 95°C for 10 min followed by 40 cycles, 95 °C for 15 sec, and 60 °C for 1 min. The quantitative real-time PCR data were analyzed by the 2-ΔΔCT method. The expression of each gene was normalized to GAPDH.

### Cytosolic Ca^2+^ measurements

An increase in intracellular Ca^2+^ was measured using the Ca^2+^-sensitive fluorescent dye Fura 2-AM as previously described (*18, 40*). The intracellular Ca^2+^ concentration ([Ca2+]i) was calculated using Invitrogen Fura-2 Ca^2+^ Imaging Calibration kit according to the manufacturer’s protocol and the equation: [Ca^2+^]free = KdEGTA × [(R − Rmin)/(Rmax-R)] × [F380max/F380min]. Kd represents the dissociation constant of Fura2–Ca^2+^ interaction, R is fluorescence ratio, R min is the ratio at zero free calcium, Rmax is the ratio at saturation calcium (39 µM), F380max is the fluorescence intensity with excitation at 380 nm, for zero free Ca^2+^; and F380 min is the fluorescence intensity at saturating free Ca^2+^. For TRPC6 peptide experiments, ECs were first incubated with 1μM of TRPC6 or control peptide for 1.5h, then loaded with Fura 2-AM fluorescent dye as described above.

### Analysis of TRPC6-mediated cationic currents

Whole-cell patch-clamp technique was used to determine cationic currents induced by WT or mutated TRPC6 channel (*10, 16, 61*). The currents were acquired during 300-ms voltage ramps from −100 to +100 mV, with a 2-s inter-ramp interval using the Optopatch amplifier controlled by PCLAMP 10 software (Molecular Devices). The current amplitudes at −90 mV were plotted as a function of time. The pipette solution contained (in mM): 10 EGTA; 3.77 CaCl2; 2 MgCl2; 125 CsMeSO3; and 10 Hepes. The standard extracellular solution contained (in mM): 145 NaCl; 1.2 CaCl2; 1 MgCl2; 2.5 KCl; 10 Hepes; and 5.5 Glucose. The pH of all solutions was adjusted to 7.2. The PCLAMP 10 software package (Molecular Devices) was used for data analysis. All electrophysiological experiments were performed at room temperature (22–23°C).

### Western blotting

ECs were lysed using radioimmunoprecipitation assay buffer (RIPA buffer containing 10 mM Tris-HCl pH 8.0, 1 mM EDTA, 0.5 mM EGTA, 1% Triton X-100, 0.1% sodium deoxycholate, 0.1% SDS, 140 mM NaCl and 1 mM phenylmethylsulfonyl fluoride (PMSF) and western blotted using indicated antibodies as described previously (*40, 62*) .

### Biotinylation assay

ECs expressing indicated cDNAs were washed with ice-cold PBS, after which cells were labeled with 0.5 mg/ml sulfo-NHS-SS Biotin in Ca^2+^/Mg^2+^ containing PBS for 30 min at 4°C. The reaction was then quenched using 100 mM glycine for 20 min, after which cells were harvested in RIPA buffer containing 10% lubrol. Equal amounts of protein were incubated with streptavidin agarose resin beads at 4°C overnight followed by three times centrifugation at 2,400 g and rinsing using RIPA buffer at 4°C. Proteins were then eluted using 4× Laemmli buffer and Western blotted as described previously (Anwar et a, 2021).

### Protein Expression and Reductive Methylation NMR analysis

TRPC6 with and without the I^111^L mutation was expressed in the BL21DE3 strain of E. coli, purified by affinity chromatography, and subjected to reductive methylation of lysine’s with ^13^C-enriched formaldehyde as previously described. Briefly, the protein at 1 mL was subjected to three successive incubations (two for 2 h and one overnight) with stirring at 4 °C. The first and the second incubations included additions of 20 µL 1 M borane–ammonia complex (NH_3_.BH_3_) (Sigma) and 40 µL 1 M ^13^C formaldehyde (Cambridge Isotope Laboratories, Inc). The final addition was 10 μL of 1M borane–ammonia complex. Glycine at the final concentration of 200 mM was necessary to end the reductive methylation reaction. The reaction biproducts were removed by buffer exchange against PBS. Next, we collected ^13^C HSQC spectra on a 600 MHz Bruker NMR spectrometer equipped with a cryogenic probe. The data was processed in NMRPipe and the chemical shifts of ^13^C methyl groups on lysines were compared for wild type and mutant TRPC6 in NMRDraw.

### Trans-endothelial electrical resistance (TEER)

HPAEC or HUVECs were seeded on gelatin-coated gold-plated eight-well electrodes (8W standard Array; 8W10E+) (Applied Biosciences, Carlsbad, CA). Cells were transfected with indicated siRNAs for 48 h. The smaller and larger counter electrodes were connected to a phase-sensitive lock-in amplifier to monitor the voltage. A constant current of 1 μA was supplied by a 1 V, 4000 Hz AC signal connected serially to a 1 MΩ resistor between the smaller electrode and the larger counter electrode. EC monolayers were incubated in a serum-free medium for 2 h, following which S1P was added to assess dynamic change in TEER (*40, 62*).

### Liposome-mediated *in-vivo* gene delivery

Liposomes were prepared using a mixture of chloroform, Dimethyl Dioctadecyl Ammonium Bromide, and cholesterol, as described previously (*16*). Briefly, chloroform was evaporated from the mixture using a rotavapor system at 105 rpm for 15-20 min at 37°C to make a lipid layer. The lipid layer was extracted by sonicating the solution for 1 h at 42°C in the presence of 5% glucose. The liposomes were filtered through a 0.45-micron filter, and cDNA was subsequently added. The cDNA-loaded liposomes were administered intravenously into the mouse. The lungs were harvested for edema measurement after 24 h. **Assessment of lung vascular injury**: The lung wet-dry weight ratio was determined to quantify lung vascular injury as described previously (*16*). Inflammatory score is assessed based on distended alveoli, capillary leakage, septal thickening, and hemorrhagic phenotype. The scoring is done manually by grading the severity of inflammation in lung histology.

### TRPC6 peptide *in vivo* studies

Animals were randomly divided for their use in therapeutic or preventive model of ALI. In the therapeutic studies, both the groups were given LPS(1mg/Kg/BW) intra-tracheally followed by administration of Control (CP) or TRPC6 peptide (TP) 15 *μ*g/50 *μ*L/mouse intra-tracheally at a gap of 2-3 hr post challenge. In preventive studies, both the groups were given Control (CP) or TRPC6 peptide (TP) 15 *μ*g/50 *μ*L/mouse intra-tracheally followed by LPS administration (1mg/Kg/BW) intra-tracheally at a gap of 2-3 hr afterwards. After 14 h, mice were sacrificed under euthanasia and lungs were harvested, left lobe was taken to measure the lung wet/dry ratio. One lobe was kept for histopathological studies and some parts were taken for FACS analysis and RNA isolation.

### Statistical analysis

Data are presented as the mean ± SEM. Data were tested for normality and equal variance to confirm the appropriateness of parametric tests. Wherever only two groups were compared, statistical differences were assessed using an unpaired two-tailed Student’s t-test. Statistical significance among three groups or over was determined using one-way analysis of variance (ANOVA) followed by post hoc Tukey’s multiple comparisons test. The data were analyzed using GraphPad Prism version 8.0, and P < 0.05 indicates a statistically significant difference.

Data points show individual values along with mean ± SD. Statistical significance: *p < 0.05, ***p < 0.001, ****p < 0.0001, ns = not significant.

## Author contribution

M. Anwar and D. Mehta conceived the study. M. Anwar and D. Mehta designed the experiments. M. Anwar, V.A. Balaji Ragunathrao, R. Amin, V. Vellingiri, J. C. Joshi, M. Tauseef, M. Z. Akhter, I. Ismail, and N. Khaki, performed experiments and analyzed the data. Ibra and I Levitan performed and analyzed patch clamp studies data. K. Chronis helped in preparing and analyzing ATAC seq and RNA seq data. H. Abdelkarim and V. Gaponenko performed and analyzed NMR data. M. Anwar and D. Mehta interpreted data and wrote the manuscript.

## Acknowledgments

This work was supported by the National Institutes of Health, USA grants HL084153, PO1-HL151327, PO1-HL160469, RO1-155941 and RO1-HL165263

The authors have no conflicting financial interests.

---

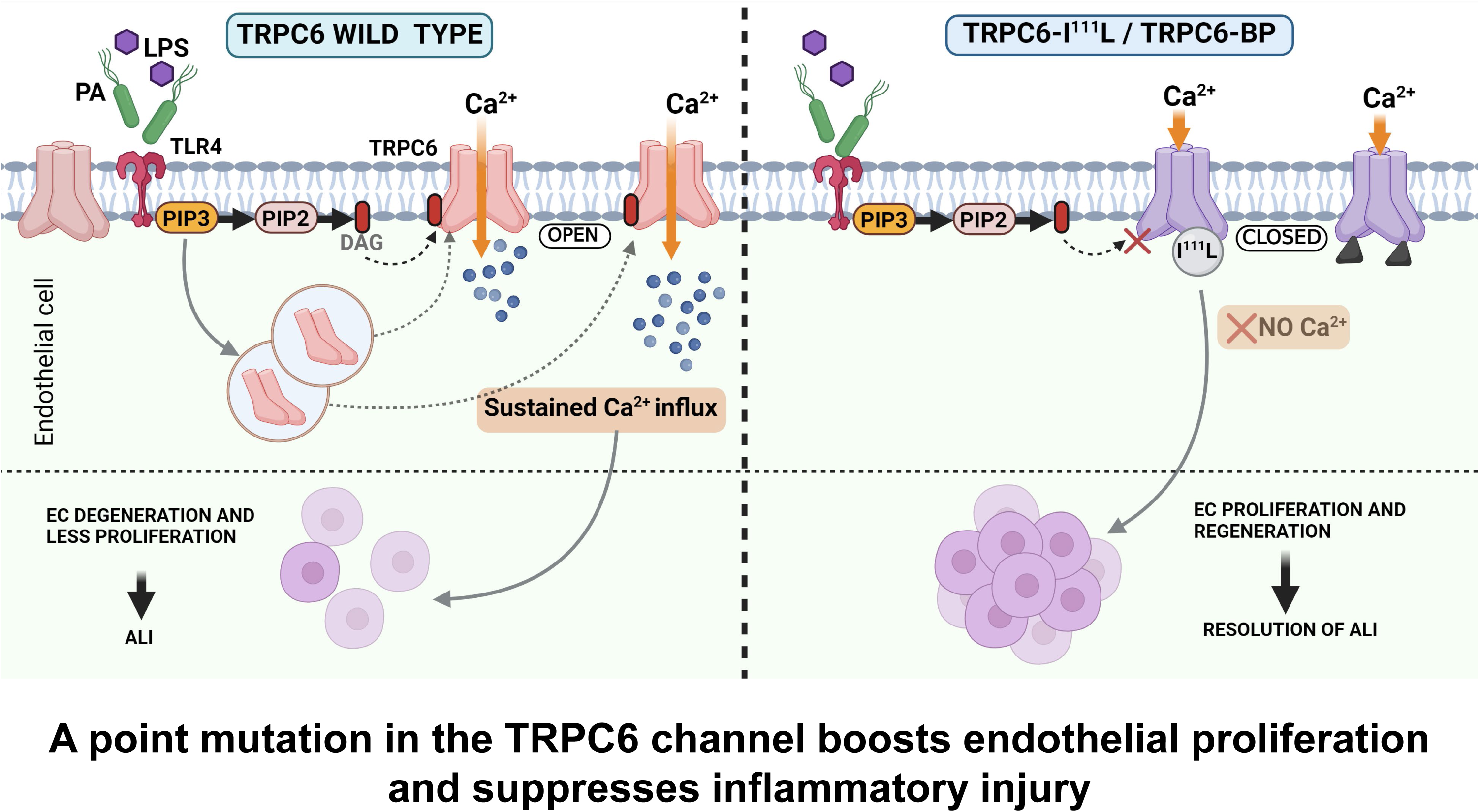

## Notes

### Competing Interest Statement

The authors have declared no competing interest.

