## Supplementary figures and images for "A Single Mutation in TRPC6 Protects Mice from Acute Lung Injury by Regenerating Endothelium"

### Supplementary Figures

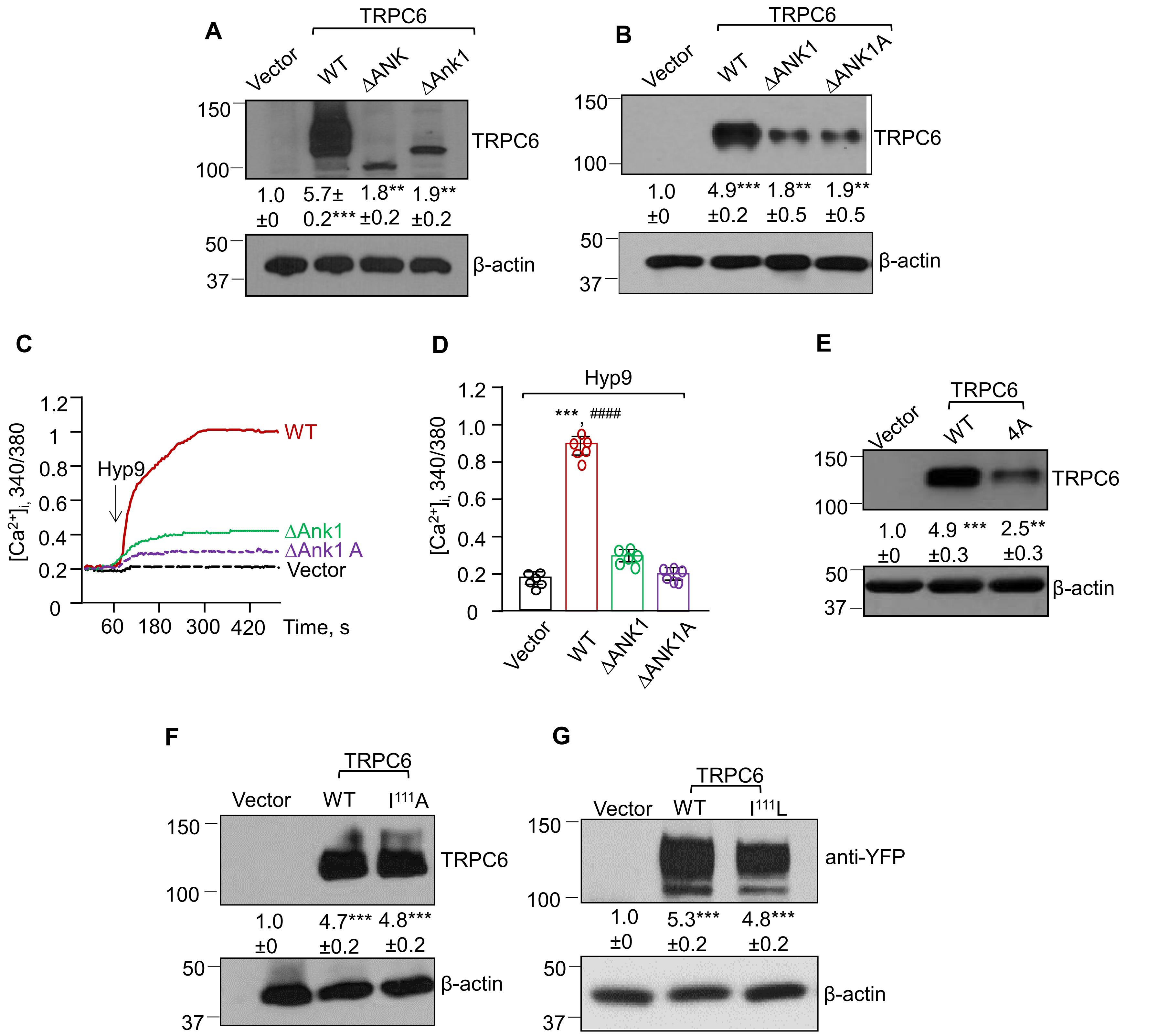

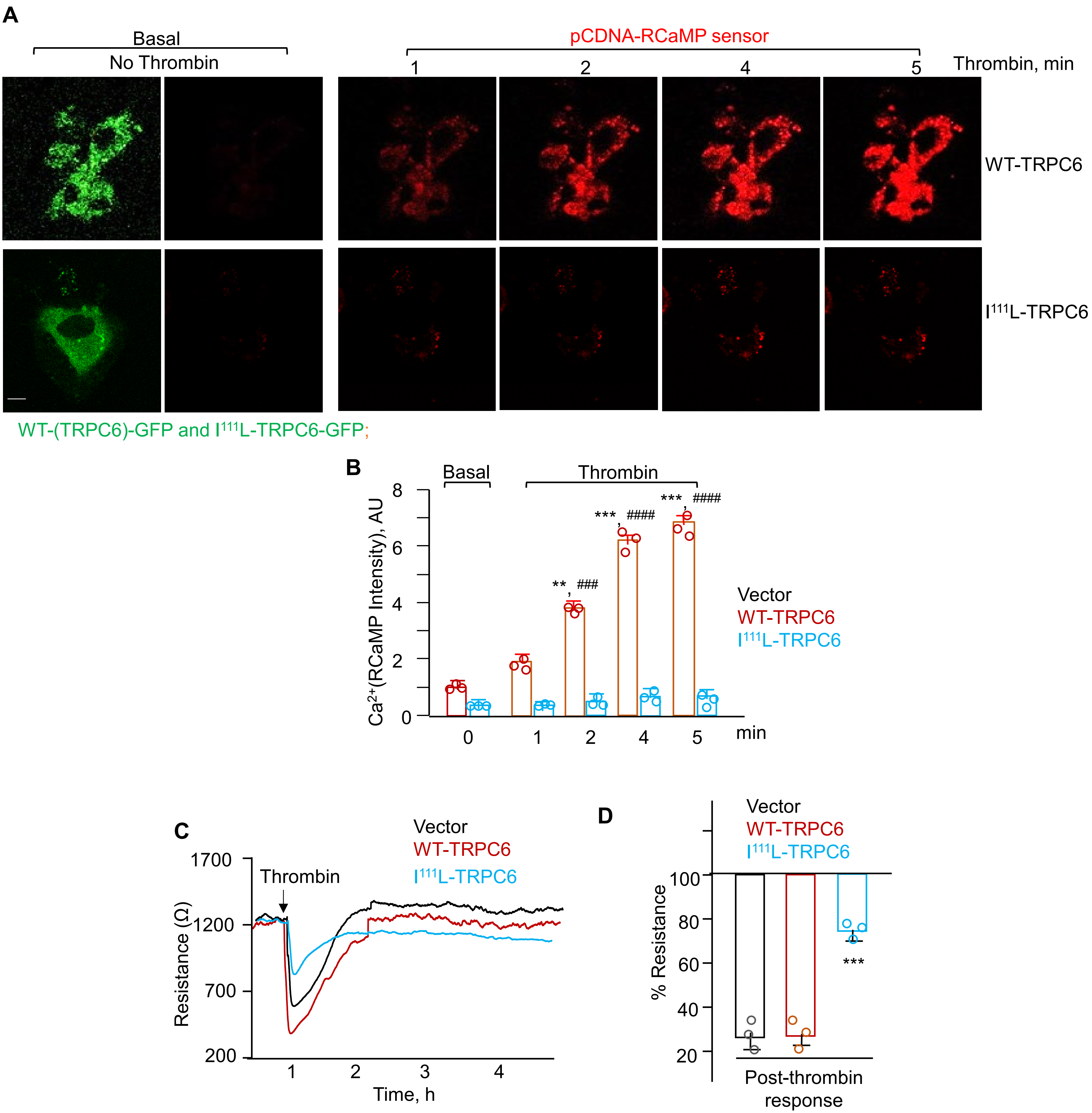

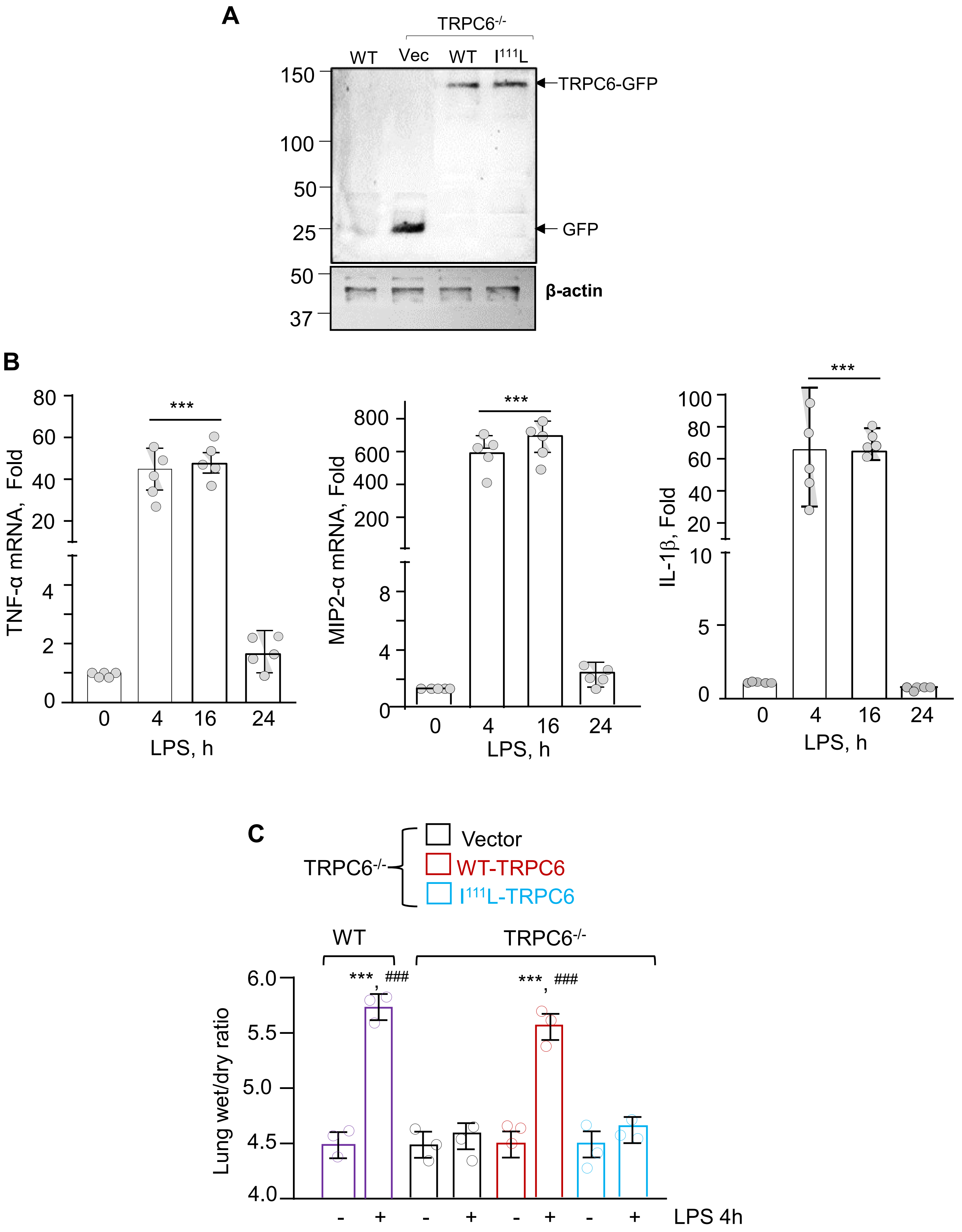

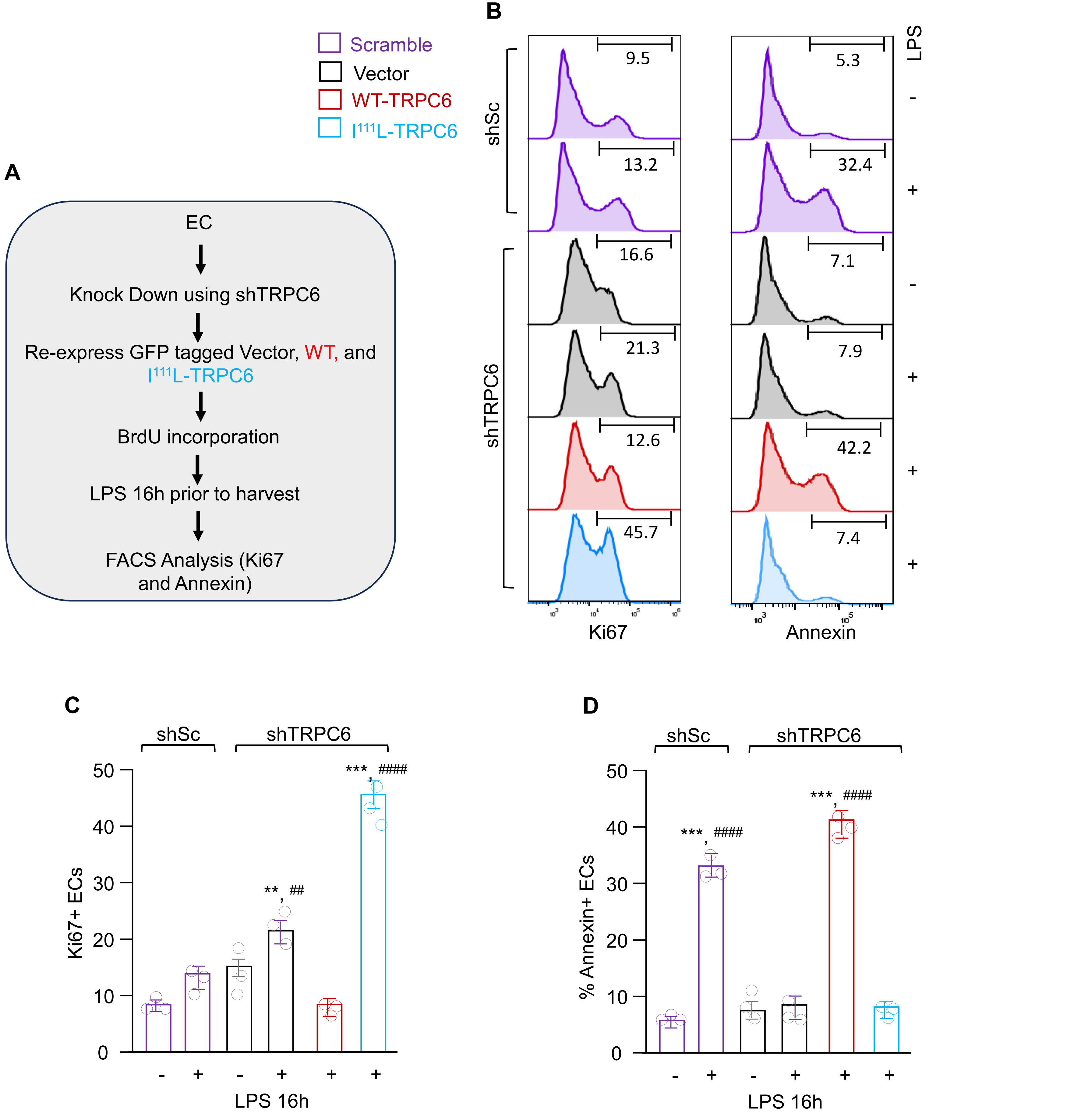

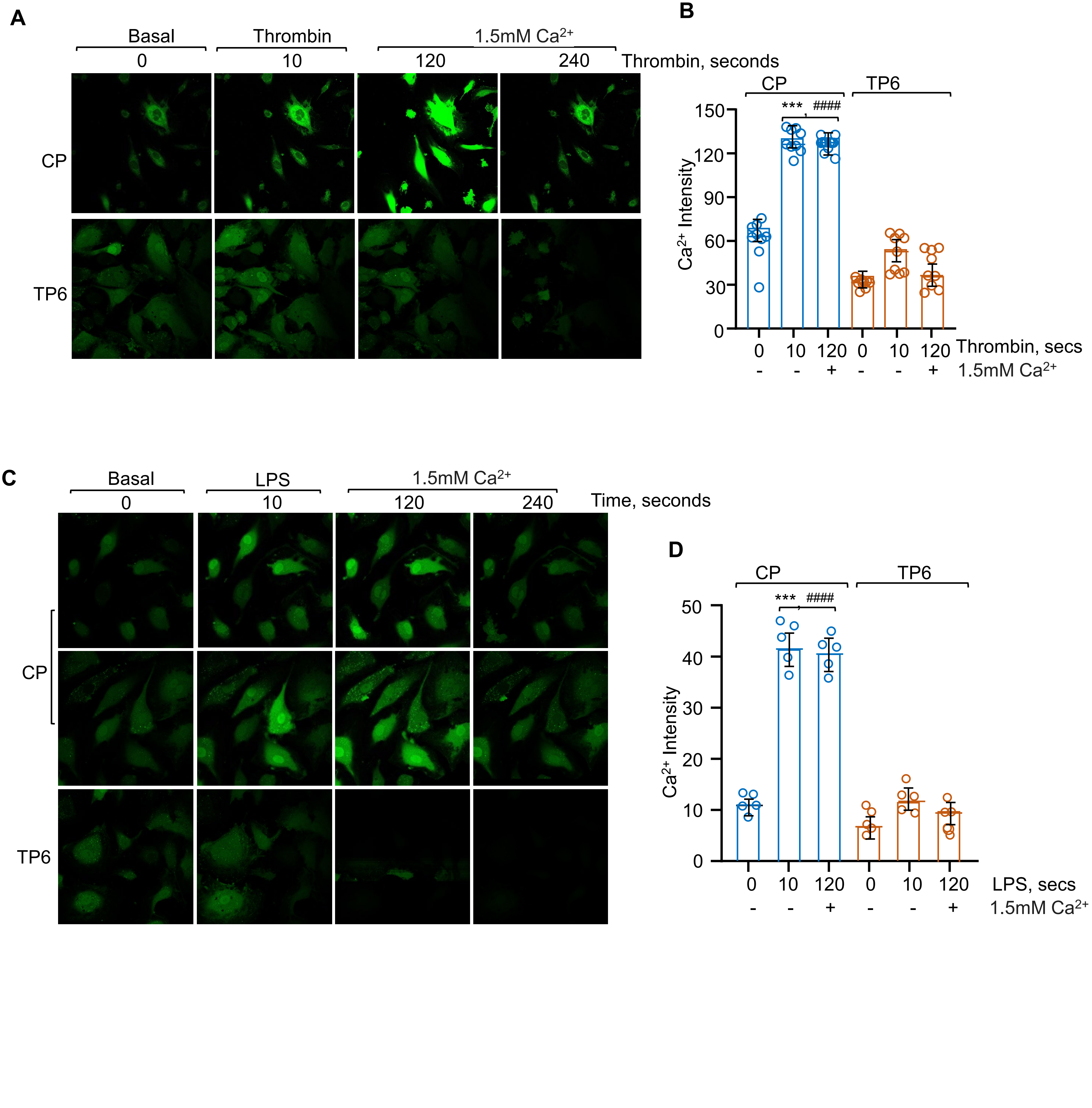

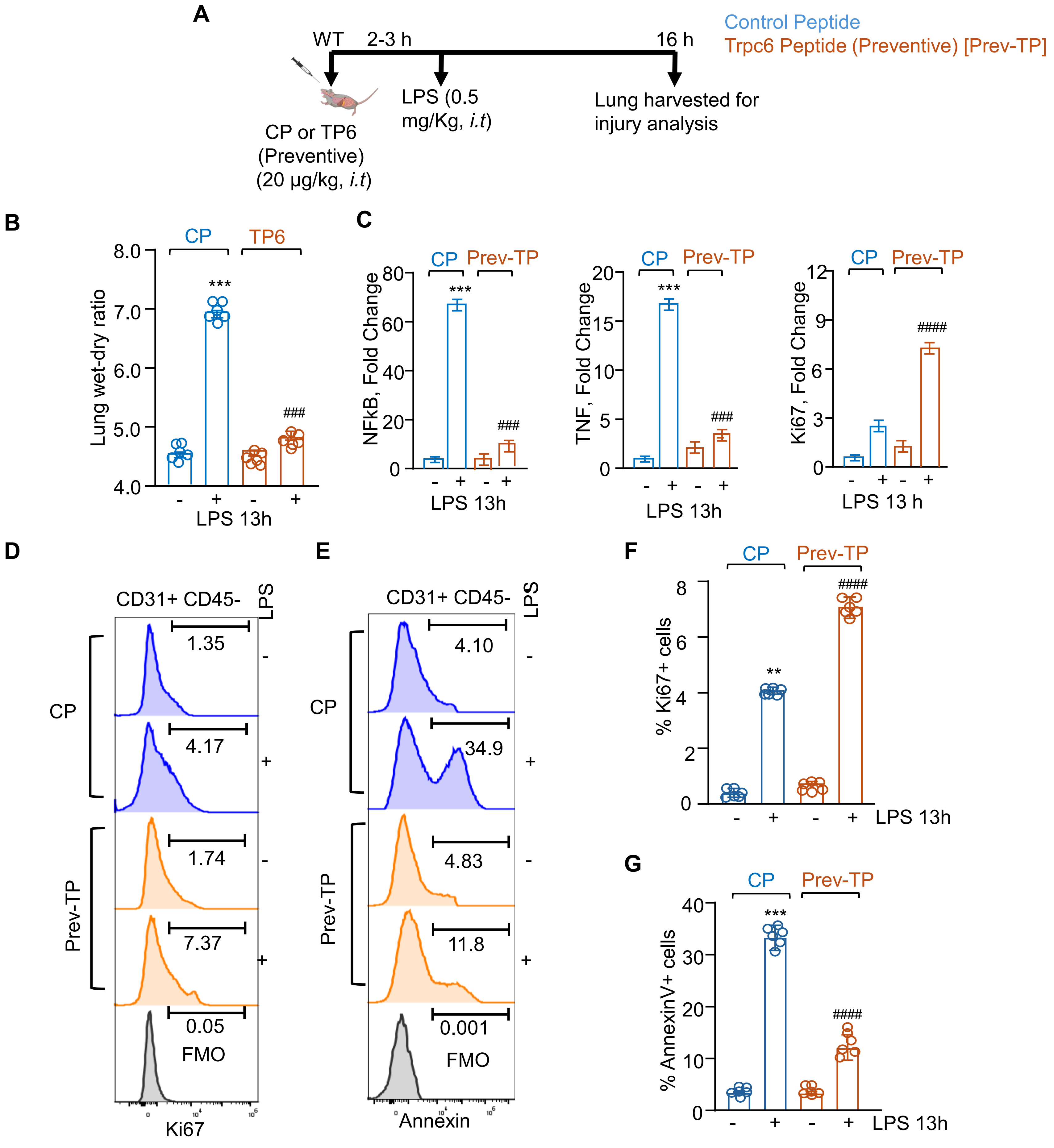
